# Flapjack: a data management and analysis tool for genetic circuit characterization

**DOI:** 10.1101/2020.10.30.362244

**Authors:** Guillermo Yáñez Feliú, Benjamín Earle Gómez, Verner Codoceo Berrocal, Macarena Muñoz Silva, Isaac N. Nuñez, Tamara F. Matute, Anibal Arce Medina, Gonzalo Vidal, Carlos Vidal Céspedes, Jonathan Dahlin, Fernán Federici, Timothy J. Rudge

## Abstract

Characterization is fundamental to the design, build, test, learn (DBTL) cycle for engineering synthetic genetic circuits. Components must be described in such a way as to account for their behavior in a range of contexts. Measurements and associated metadata, including part composition, constitute the test phase of the DBTL cycle. These data may consist of measurements of thousands of circuits, measured in hundreds of conditions, in multiple assays potentially performed in different labs and using different techniques. In order to inform the learn phase this large volume of data must be filtered, collated, and analyzed. Characterization consists of using this data to parameterize models of component function in different contexts, and combining them to predict behaviors of novel circuits. Tools to store, organize, share, and analyze large volumes of measurement and metadata are therefore essential to linking the test phase to the build and learn phases, closing the loop of the DBTL cycle. Here we present such a system, implemented as a web app with a backend data registry and analysis engine. An interactive frontend provides powerful querying, plotting and analysis tools, and we provide a REST API and Python package for full integration with external build and learn software. All measurements are associated to circuit part composition via SBOL. We demonstrate our tool by characterizing a range of genetic components and circuits according to composition and context.

## Introduction

Synthetic biology is an interdisciplinary field that aims to use engineering principles to create novel DNAs with prescribed functions. This process typically follows a design-build-test-learn (DBTL) cycle (figure 1). DNA design has benefited from developments in syntax,^1–3^ part registries,^4,5^ BioCad tools,^6,7^ standards for describing and sharing designs^8^ and web-based information systems.^9,10^ These tools enable exchange between a collection of open source tools^11^ that implement phases of this engineering cycle (figure 1). For example, simulation tools^12,13^ can be used to predict candidate genetic circuits, from which potentially functional circuits are selected. All circuit compositions and part sequences can be designed using the above mentioned tools to generate files in different formats such as XML, GenBank, FASTA or SBOL. These designs could be stored and published on SynBioHub^14^ or on JBEI-ICE registry,^5^ receiving a unique identifier making possible for the constructs to be accessed by specialized software tools.

**Figure 1:**
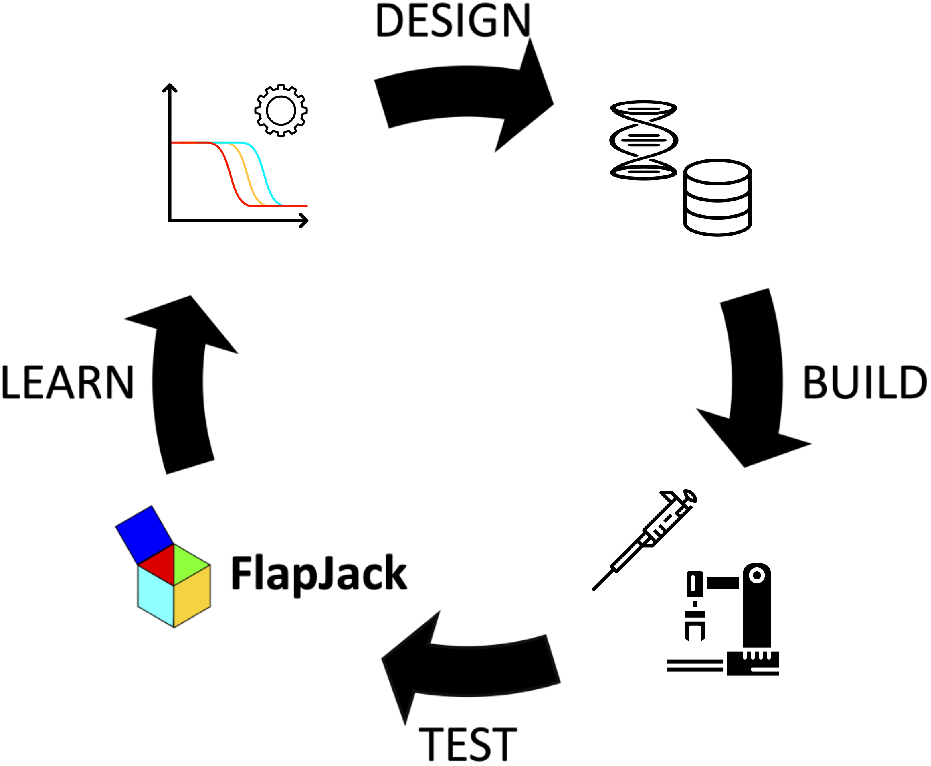
DBTL cycle. The design-build-test-learn (DBTL) cycle can be mostly implemented with existing open source software tools. Flapjack closes the gap between test and learn phases, organising and analysing measurements to parameterize or train models.

The DNA sequence designs generated are then built using modern assembly techniques.^15–18^ These techniques are increasingly enabled by robotic automation,^19–21^ controlled by open source application programming interfaces (APIs) that can be integrated with different standards. Robotic automation can also be combined with scaleable low cost and open source measurement devices that can be run in parallel to generate high-throughput test data.^22,23^ These test data feed into the learn phase to estimate parameters of models,^24^ train machine learning algorithms,^25^ or define design constraints.^26^

Automation is rapidly expanding our capacity to build and test large numbers of genetic circuit designs for applications from microbes^27^ to higher organisms such as mammals.^28^ However, there remains a need for tools to link these large volumes of test phase data to methods and algorithms to learn new designs, closing the DBTL cycle (figure 1). Fundamental to the DBTL cycle is characterization of component (part, device, circuit) function based on test data. This data must be collated, filtered and analyzed to provide the required inputs for the learn phase.^26^ Metadata not present in raw data files is also necessary to incorporate details of the DNA design (part composition), experimental conditions (temperature, media etc.) and chassis^29^ in which measurements were made. Combining all of this information into a coherent predictive model is the goal of the learn phase.^30–32^

Genetic circuits are dynamical systems driven either by a host cell^30^ or a cell-free substrate.^33^ To fully characterize their behaviour it is therefore necessary to measure and analyze the dynamics of their gene expression. There is currently no standard for the required kinetic measurement data, which is distributed across many files in different formats, stored in different labs, institutions, repositories and databases. While excellent systems biology analysis tools exist for studying gene expression dynamics^34–36^ these operate on single experiments, do not incorporate metadata nor DNA part composition, and do not integrate with searchable registry databases. Neither do existing measurement data registries^10,37–39^ reference DNA design part composition via exchange standards, such as SBOL.

Here we present a tool which completes the DBTL workflow shown in figure 1A; storing, filtering, visualising and analysing kinetic gene expression data from the test phase in order to provide input to the learn phase (figure 1A).

## Results

### Data model

Central to our system is the data model, which defines how the measurement data is organized and labelled (figure 2, table 1). The basic unit of data is the *measurement*, which typically consists of readout of fluorescent, luminescent or colorimetric reporter level,^40,41^ and estimates of biomass or density. Each measurement refers to a particular *sample* at a specific time, and each sample is measured at multiple time points, possibly reading multiple reporter levels. Each sample is measured in a specific *assay*, which might correspond for example to a kinetic plate-reader experiment. Finally, sets of assays are grouped into a *study* representing a project with a common purpose. This hierarchy represents typical lab workflows and allows users to easily filter data intuitively, providing a platform for data exploration.

**Figure 2:**
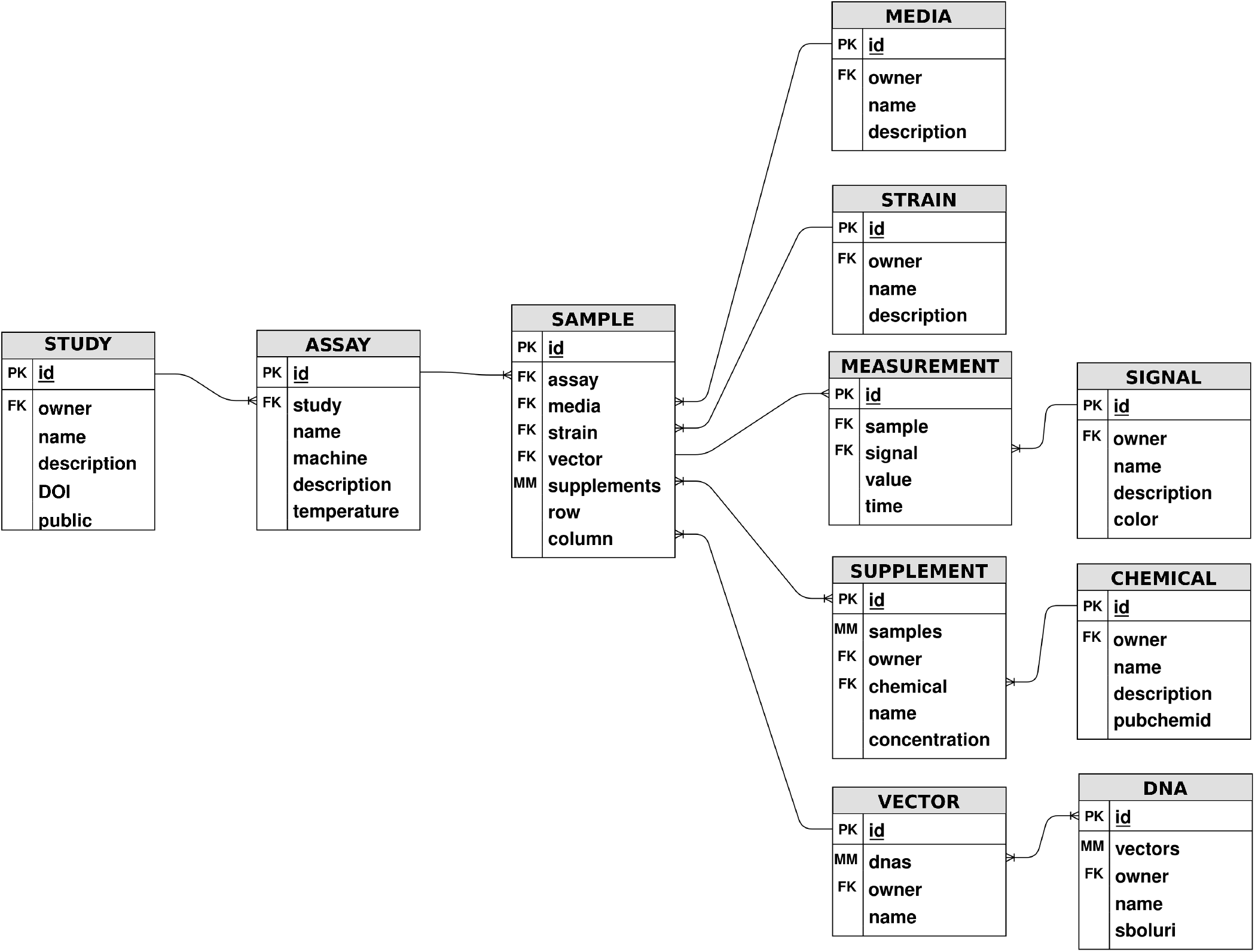
Flapjack’s data model. The basic unit of data is the *measurement*, which is the readout of a reporter or biomass *signal* in time. A *measurement* refers to a specific *sample*, which can be a well in a plate-reader experiment or a colony in a petri dish time-lapse. *Samples* are grouped in a specific *assay*, that corresponds to a single kinetic procedure. *Assays* belong to a particular *study* that aims to answer a scientific question or project. Along with *signal* data, each *measurement* is described in terms of its context through experimental metadata, mainly composed of the *media* substrate, the host *strain*, the *DNA* which the host is transformed with and inducer *chemicals* to study stress or other regulation behaviours of the system. It is important to mention that some metadata could not be present, such as *strain* in Cell-free *assays* or *DNA* in control *samples*, to name a few.

**Table 1:**
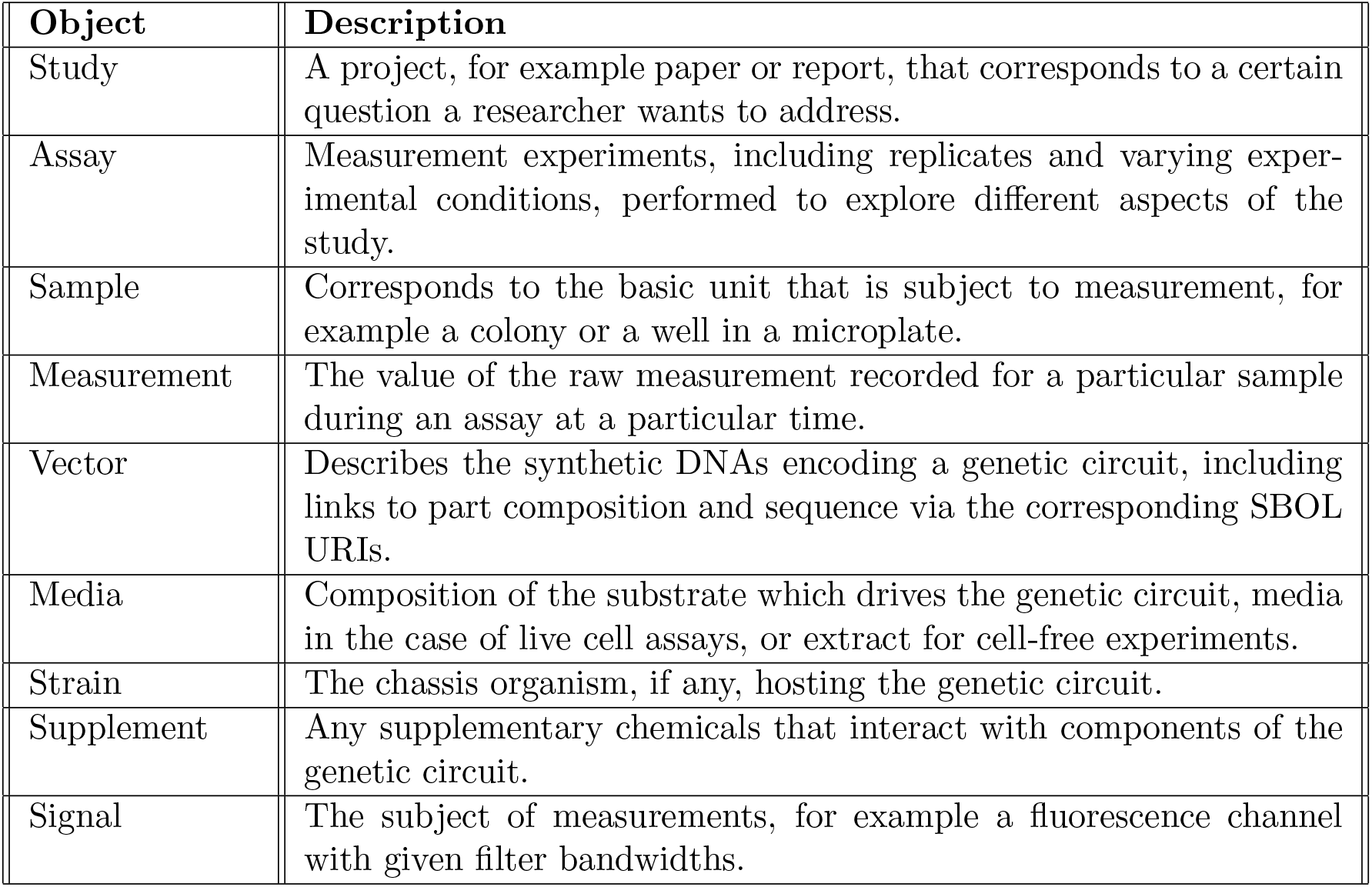
The objects that constitute the Flapjack data model.

However, in order to fully characterize the behaviour of genetic circuits and their components one must consider the context^29,42^ in which measurements were recorded. This context is given by metadata which are stored in our database and related to each measurement. For example, to account for metabolic changes, the chassis strain used, the media in which it grew, and the assay temperature must be considered during data analysis. External signals in the form of inducer chemicals must also be incorporated to parameterize models of gene regulation. Most importantly, using the stored SBOL URI of genetic circuits and connecting to SynBioHubs^14^ *via* PySBOL,^43^ we can analyze the composition of the DNA and relate the context of components to their effect on circuit behaviour. This combination of measurement data and contextual metadata enables sophisticated analysis and modelling^30^ of genetic circuits that can be used to learn increasingly complex designs^44,45^. Identifying DNAs by their SBOL URI means that Flapjack effectively integrates measurement data, its analysis, and its metadata to the exact sequence of parts from which the circuit DNA is composed.

### Web application and API

The architecture of Flapjack is shown in Figure 3 and it is composed of a web interface frontend and a backend application programming interface (API). The connection between frontend and backend is made via a REST API provided by the backend and WebSocket connections for functionalities that require a persistent connection. Both the API and WebSockets function with authentication via JSON (Javascript object notation) web tokens, which are required to access restricted endpoints. JSON is used for all data exchange, maximizing flexibility and interoperability with external tools.

**Figure 3:**
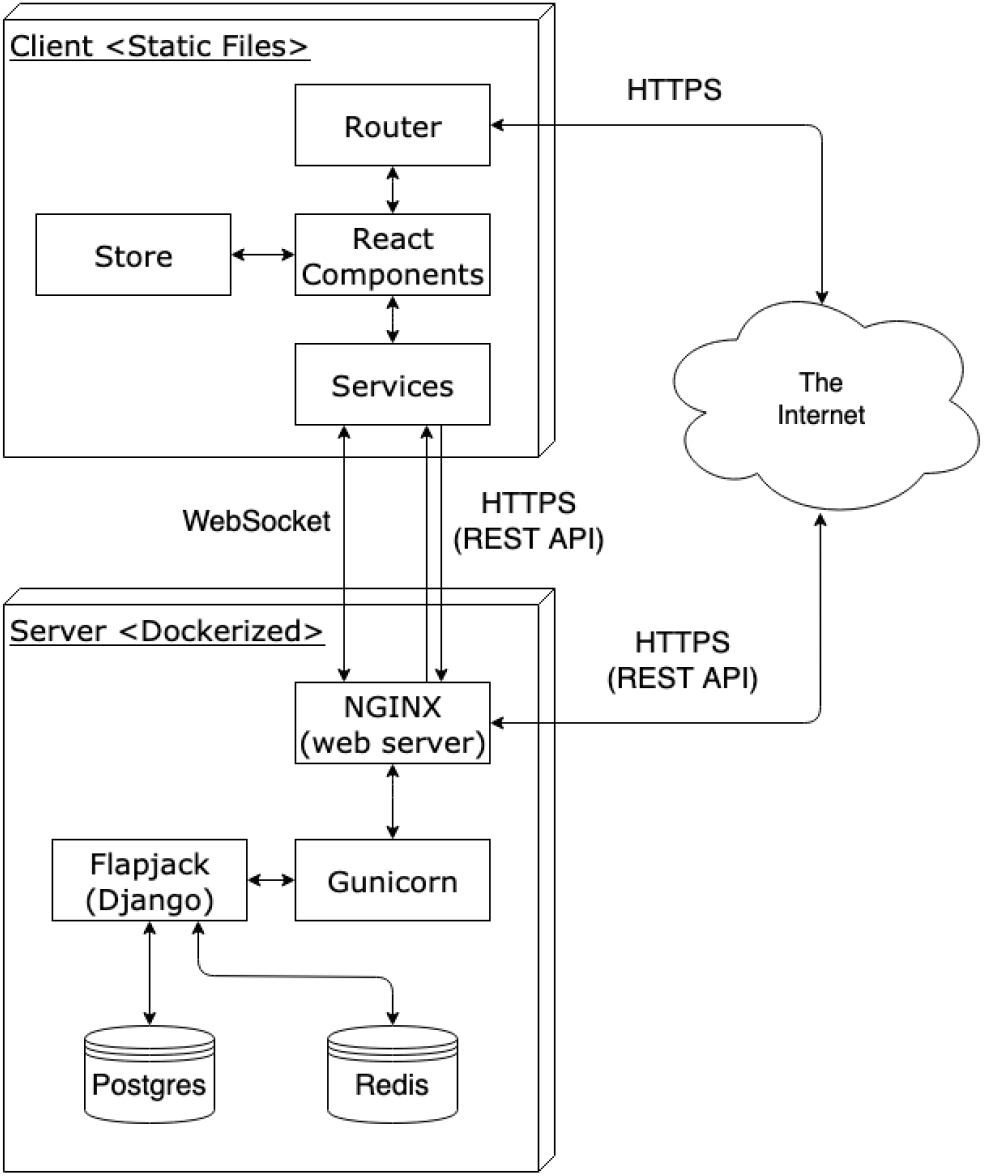
Flapjack’s architecture. Flapjack is a full stack web application developed as a Django REST API and a React web client, built on a PostgreSQL database and deployed in Docker containers for ease of use and portability. The frontend-backend connection is made via the REST API and WebSockets when persistence is needed, both using JSON web tokens for authentication to access restricted endpoints. The backend makes use of NGINX to reverse proxy to a Gunicorn application, Redis database to store data related to WebSockets and data science libraries to process data and perform analysis such as NumPy, Pandas and SciPy. The frontend uses D3.js library via Plotly to visualize data, Redux and Redux Persist for global state management and data persistence. We also provide a Python package to access progammatically to the REST API. In order to facilitate interoperability with external tools, all data exchange is made using JSON.

The backend is written in Python,^46^ using the Django framework as a base for the API and WebSockets architecture. The Django Rest Framework library is used to build a powerful and flexible REST API, while Pandas,^47^ Numpy^48^ and SciPy^49^ libraries are used to process the data before sending it to the frontend. The API provided by the backend, in addition to serve the frontend with the services it needs, can be used to interact with Flapjack for more sophisticated analysis by connecting to existing libraries and software^14,47,50–52^ through a provided Python package^53^ (tutorials for the package are also provided^54^). In order to achieve performance and security in the backend, an NGINX server is used to reverse proxy to a Gunicorn application that serves the Django Application, and to process the requests. All persistent data is stored in a PostgreSQL database, while a Redis database is used to store data related to WebSocket connections and messages. The backend is deployed using Docker^55^ for easy installation and portability.

The frontend interface is written in JavaScript, using the React library^56^ for an interactive design, the D3^57^ visualization library via Plotly^58^ and Redux for global state management and data persistence. This application is intended for convenient data exploration and basic analysis, and can also be deployed using Docker^55^ for ease of setup. Access to user data in the frontend is managed via JSON web tokens, where the refresh token is stored in the browser’s local storage via Redux Persist, and is used upon site load to obtain the user’s access token. In the first instance, the user (owner) who uploads the data is the only one who has access to it. We have implemented a data sharing system that considers giving access to particular users or making studies public. Thus, we ensure that only those authorized by the owner have access to the information.

### Web interface

We provide a comprehensive web interface that firstly allows users to create a new study, or add assays to an existing study, by uploading data files and inputing metadata interactively *via* the web app. The information is provided to the platform through a modified version of the excel files exerted by microplate readers or a JSON file in the case of FluoPi assays. Information about how to format these files can be found in out tutorials.^59^ On the browse page, the user may browse and search available studies, assays and vectors. Here the user may modify studies to which they are owners; deleting, sharing, or publishing. On a separate tab users may also search and browse a table of assays, viewing a short description of each. The vector tab provides an optional SBOL URI which links to a SynBioHub instance^60^ where it is possible to download the SBOL design file^14^ and SBOL circuit diagram,^61^ as well as inspect part composition.

On the view page (figure 4) Flapjack provides a tabbed environment in which to explore and analyse data. Users construct a query by selecting filters for study, assay, media, strain, vector, and signal. For example, one might query all data for a given vector, irrespective of the study in which it was performed. This data can then be visualised interactively, plotting each measurement as a function of time. Plots are grouped into sub-plots, and given different line colors according to user-selected labels. For example the user might compare two studies by creating two tabs in the user interface. Within each tab the user could also select to group the data into sub-plots according to the vector in each sample. Finally it would be possible to group the lines into colors according to the signal, for example to differentiate fluorescence channels. Data can also be summarized by plotting mean and standard deviation and/or normalized for comparison. The plots generated by the frontend can be downloaded as print quality PNG files with specified dimensions and font size, or as JSON encoded figure objects for formatting with Plotly.^58^ Plots are stored in the browser’s local storage for persistence while sessions are active.

**Figure 4:**
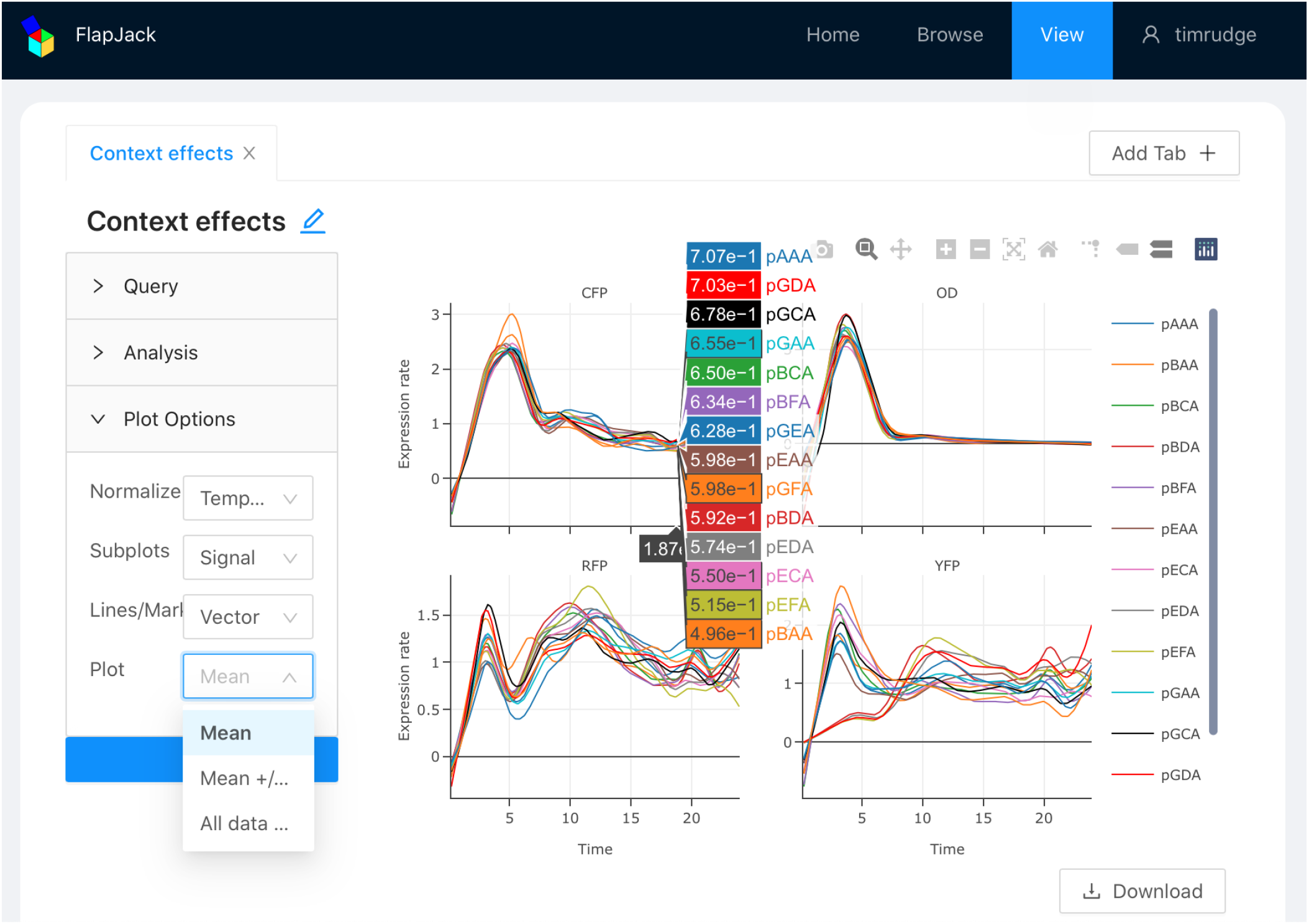
The web interface of Flapjack. The view page allows users to query data stored in the database and perform a variety of analysis. Queries and analysis can then be visualised interactively in plots grouped into sub-plots and given different line colors, both according to user-selected labels (i.e. *assay*, *vector*, *signal*, etc). Also, data can be summarized by plotting mean and standard deviation and/or normalized for comparison. These plots can be organized in different tabs and stored in the browser’s local storage while the session is active, thus users can compare between different querying filters. Finally, users can download the generated plots as PNG files or as JSON encoded figures for formatting with Plotly and use them for publication purposes.

Having constructed a query the user can also perform various analyses, plotting the results according to the same groupings. The user provides input parameters to a range of functions that transform the selected data time series. For example, the *mean expression* summarizes a kinetic experiment by the average reporter level, and results in a set of bar graphs according to the specified grouping options. Flapjack automatically detects control samples, those without cells (media background) and those without DNA (untransformed cells), and computes and subtracts signal background where necessary. More complex analyses include calculation of expression (or synthesis) rates, induction curves, velocity profiles, and induction kymographs. Growth can also be considered, using the same tools to extract mean or maximum biomass or growth rate by analyzing measurements that correspond to biomass estimates. A comprehensive set of tutorials for the capabilities and use of the web interface is provided on our wiki.^59^

All querying, plotting and analysis functions and resulting figures and data may also be accessed *via* our Python package^53^ (tutorials provided^54^). Via the API, this package interfaces Flapjack directly with Pandas^47^ and thus to the Numpy/SciPy stack^48,49^ and various machine learning packages.^50,51^ Figures downloaded as JSON may be formatted and added to (e.g. annotated) using Plotly^58^ via Python^46^ or JavaScript. Our API uses JSON making it simple to interface with other software packages.

### Design-build-test-learn cycle

We used Flapjack to complete the DBTL cycle (figure 1), which can be summarized as follows. First, in the design phase circuits were composed using SBOLDesigner^7^ and uploaded to our SynBioHub instance.^60^ The build phase, DNA assembly, was performed manually but could be automated using open source APIs that can be linked to genetic designs.^16,21^ Test phase data was generated using various kinetic gene expression measurement techniques, which can be automated to varying degrees.^20–23^ Test data and metadata was then uploaded to Flapjack, allowing visualization and analysis of the synthetic genetic circuits measured under various conditions. This data was then used to quantitatively characterize circuit behaviour, allowing estimation of parameters for models to predict new designs, which could be simulated using custom software or existing simulation tools.^12,13^ These simulations can predict novel circuit behaviour, constituting the learn phase.^30^ Furthermore, Flapjack provides *tidy* analysis data with characteristic features linked to metadata labels that can directly feed into machine learning and statistical analysis toolkits.^50,51^

In the following sections we demonstrate the functionality of flapjack by characterizing a range of genetic circuits in a range of contexts.

### Characterizing context effects on gene expression levels

Gene expression can be strongly context dependent,^29,42^ depending on neighbouring genes (compositional context), host strain, and media composition (cellular context). To examine these effects we designed and assembled 14 combinatorial three-reporter plasmids, combining 10 different transcriptional units (TUs) which were driven by one of seven promoters.^62^ Three of these promoters were constitutive synthetic promoters,^1^ two were repressed by endogenous repressors LacI and TetR produced by strain MG1655z1, and two were inducible but measured in the absence of inducers to study their basal expression. Each plasmid contained three transcription units producing RFP, YFP and CFP. The CFP TU was maintained the same in all plasmids, to serve as a reference.^63,64^ Plasmids were named by letters representing each of their three promoters (see Supplementary Methods for details of parts). The down-stream RBS, CDS and terminator for each reporter was maintained constant. This collection of plasmids (figure 5A) presented each TU in multiple compositional contexts, that is, in the presence of different up-stream and down-stream genetic parts.

**Figure 5:**
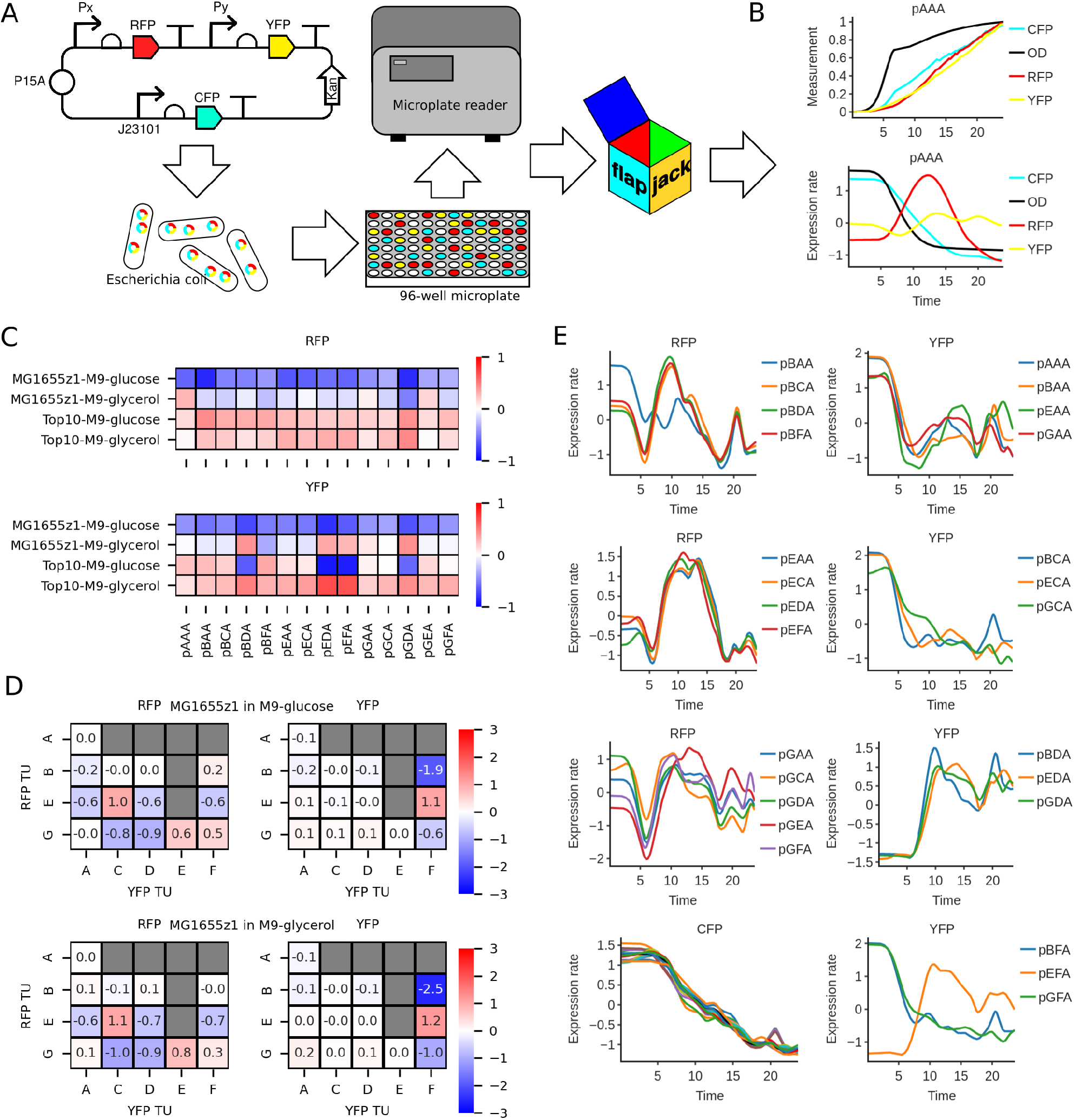
Characterizing context effects on gene expression levels. (A) Three-reporter plasmids containing different TUs to study compositional context and in different strains and media to study cellular context. (B) Once data is uploaded to Flapjack, users can ensure data quality before proceeding with more complex analysis. (C) Cellular context: mean fluorescence levels of RFP and YFP, red being above and blue below mean. Relative magnitude of gene expression was affected by changing media and strain. (D) Compositional context: mean fluorescence levels of RFP and YFP depending on up-stream and down-stream genetic parts. RFP is more affected by down-stream YFP than YFP is by up-stream RFP. (E) Expression rate for RFP and YFP in all plasmids show similar profiles, considering this collection of TUs relatively orthogonal.

The plasmids were transformed into Escherichia coli, grown in 96-well plates and their fluorescence and optical density (biomass) measured in a microplate reader for 24 hours (figure 5A). The cellular context in which these genetic circuits were tested was varied by measuring them in two different strains of E. coli, and in two different carbon sources. The resulting fluorescence and optical density traces could then be analyzed using the Flapjack front-end to compute the growth rate and gene expression rates (figure 5B). Further analysis reveals the effects of compositional and cellular context on the magnitude and time dynamics of gene expression.

Flapjack allows us to easily compare the magnitude of expression from each plasmid in each of the combinations of strain and media composition (figure 5C). First we compute the mean fluorescence of each sample during the assay. In order to normalize the levels of expression to be comparable we divide by the expression level of the reference CFP TU.^63,64^ Next to understand the effect of context independently of overall TU strength, we normalize each column by the mean for that plasmid, so that red values are above, and blue values are below the mean. We can immediately see that the relative magnitude of gene expression was affected by changing media and strain. This suggests specific contextual effects on each TU rather than global metabolic regulation of gene expression, for example due to varying polymerase levels, which would equally affect the reference CFP expression. In the case of the RFP TU the fold changes are consistent for all plasmids, whereas the downstream YFP TU is much more variable in its response. This may be due to interference from the upstream TU due to transcriptional read-through^65^ or supercoiling.^42,66^

To understand the effects of compositional context, Flapjack enables us to use the SBOL URI of each plasmid to query its part composition from SynBioHub.^14^ Given the composition of RFP and YFP TUs we can then construct a similar map of fold changes in TU expression depending on up-stream and down-stream genetic parts (figure 5D). We see that the RFP is subject to variation due to the composition with the down-stream YFP TU. The YFP TU is less affected by its upstream RFP TU, except for the outlier case of inducible promoter pLas81 expressing YFP (TU “F”, see Tables 1 and 2 in Supplementary Information) which shows more than 4-fold variation depending on the TU to which it is adjacent. Despite these context effects on our collection of reporter TUs, the characterization shows that the order of magnitude of relative gene expression was conserved across all contexts meaning that this collection of TUs may be considered orthogonal for some purposes.

### Effect of context on gene expression dynamics

Expression of genes is in general not constant but varies over time. To fully characterize our collection of reporter TUs we must therefore consider their dynamics. Consider a simple one-step model of gene expression,

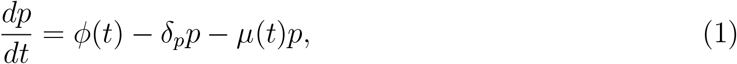

where *δ_p_* the protein degradation rate, *μ*(*t*) the time varying growth rate, and *ϕ*(*t*) is the dynamic gene expression rate profile. The growth of biomass *μ*(*t*) is defined as,

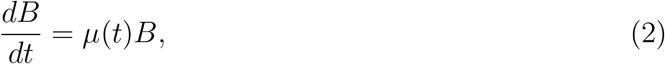

and the background corrected reporter measurements *y*(*t*) ∝ *Bp*, with *B* the background corrected optical density (OD). The expression rate profile can be computed from this system of equations by noting that for *δ_p_* = 0 (stable reporter),

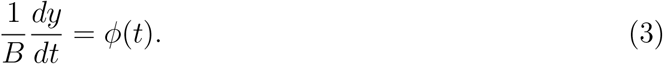

To avoid amplification of noise Flapjack computes this indirect expression rate profile by applying a Savitsky-Golay^49^ smoothing filter to the measured time series *B*(*t*) and *y*(*t*), differentiating the smoothed fluorescence intensity time series, and finally normalizing the derivative by the smoothed biomass time series.

However, this method is highly sensitive to noise. Direct inferrence of the expression rate profile using linear inversion^35^ has recently been shown to overcome this amplification of noise. Flapjack implements the linear inversion method via the WellFare Python package provided by the originators of the approach.^35^ We computed the expression rate using the direct linear inversion method for each of the combinatorial plasmids in each fluorescence channel. Based on the part composition, we can plot the expression rate profiles of each TU in its range of compositional contexts. In figure 5E the temporal mean normalized expression rate of each RFP TU is plotted in each of its compositional contexts (left column), and similarly for YFP TUs (right column). In each plot the same TU is measured in different compositional contexts. We can see that the essential dynamics of gene expression are unchanged by the compositional context, except in the same outlayer case as above - expression of YFP driven by promoter pLas81 (YFP TU “F”, see Tables 1 and 2 in Supplementary Information) in the presence of promoter the repressible promoter R0040 upstream (RFP TU “E”, see Tables 1 and 2 in Supplementary Information).

Here we show only the case of strain MG1655z1 and M9-glucose media, but the same result is repeated in all conditions (see Supplementary Information for full data).

We note that RFP and YFP TUs have distinct gene expression rate profiles. Consider the particular case of pAAA, in which each TU contains the same constitutive promoter (figure 6A). Despite being driven by the same promoter, each TU exhibits different expression dynamics, with CFP and YFP broadly following growth rate, while RFP has a later peak. To determine if this was an effect of gene composition order (RFP followed by YFP), we reordered the TUs to generate plasmid pAAAF (figure 6B). The gene expression profiles were essentially unchanged, suggesting that the RBS, CDS and fluorescent reporter itself may influence gene expression dynamics. This result shows that caution must be taken when interpreting reporter expression profiles as representative of promoter activity.

**Figure 6:**
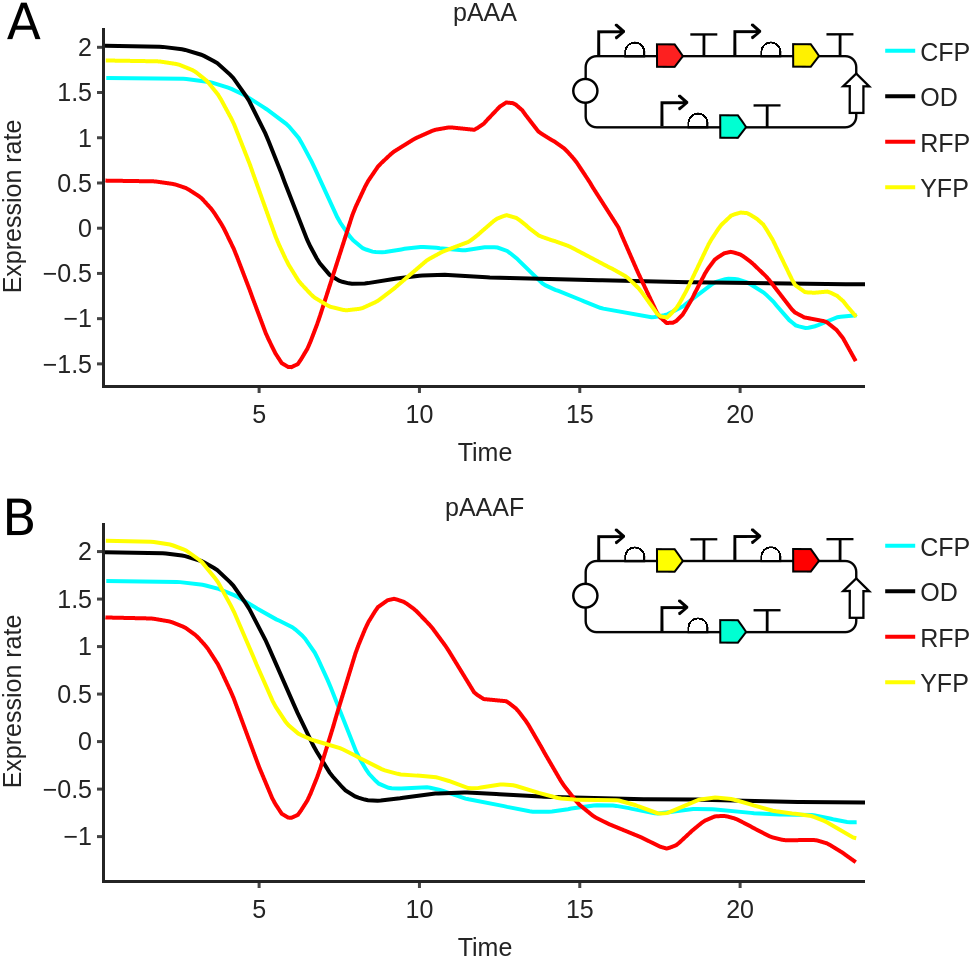
A. In the particular case of pAAA plasmid, expression rates of YFP and RFP follow growth rate while RFP has a later peak. B. RFP and YFP TUs were flipped in order to determine if a gene composition order was affecting these profiles. Gene expression profiles were essentially unchanged, suggesting that the RBS, CDS and fluorescent reporter itself may influence gene expression dynamic.

### Characterizing CRISPRi transcription regulation

Genetic circuits consist of multiple genes interconnected via regulatory interactions. This means that the product of one gene affects the expression of other genes in the circuit. Here we consider regulation by CRISPR (clustered regularly interspaced short palindromic repeat) interference.^67^ CRISPRi functions via expression of a deactivated Cas9 enzyme (dCas9) which is directed by a single guide RNA (sgRNA) to bind to a region of the target gene, preventing transcription. The efficiency of repression depends on the sequence homology of the sgRNA to the target region. Here we targeted an mVenus fluorescent reporter gene with three different sgRNA sequences. We varied the expression of dCas9 via IPTG induction and maintained expression of sgRNA at constant high expression, so that dCas9 concentrations were rate limiting (figure 7A). We assembled three modular inverters or NOT gates based on this design principle, with a common constitutive reference transcription unit (figure 7A). For the reference we used the same promoter as in the previous study^63,64^ driving mTurquoise2 fluorescent protein. These genetic circuits were measured in a microplate reader as in the previous study, in a range of IPTG concentrations. We expect that at high levels of IPTG the dCas9 will repress the expression of mVenus fluorescent protein.

**Figure 7:**
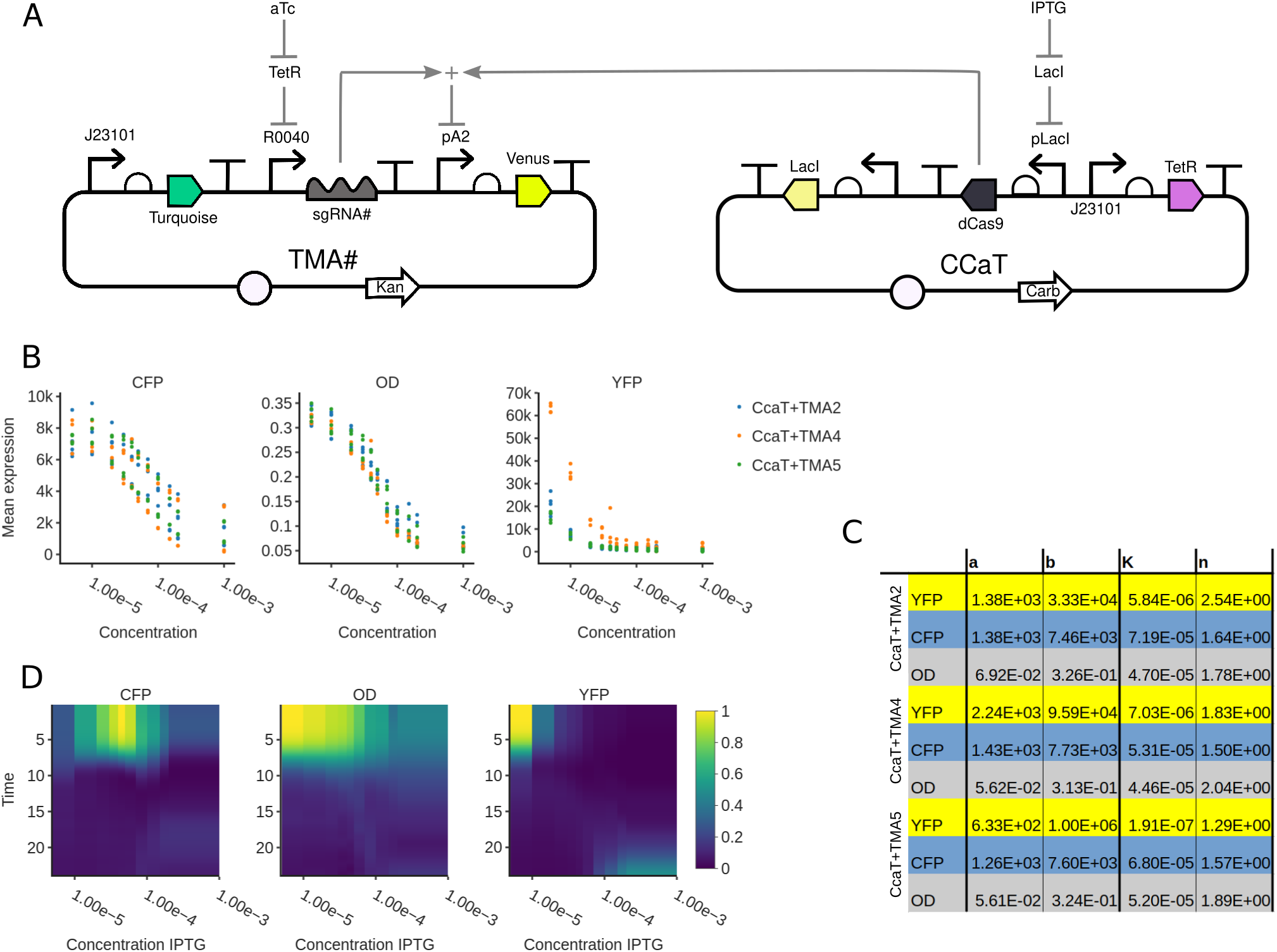
CRISPRi regulation of Venus expression. (A) Three different sgRNA sequences that direct the enzyme dCas9 to a region of the target mVenus fluorescent reporter gene, preventing its transcription. IPTG expresion was varied, expecting that at high levels the dCas9 will repress the mVenus expression. aTc regulates the expression of sgRNA and was mantained at constant high expression. (B) Transfer curves as the output of logic gates. Using Flapjack we easily plot the mean expression as a function of IPTG concentration for the biomass and reporters. Growth rate, as well as reference and target TUs were strongly repressed by the presence of dCas9. (C) Using the Flapjack Python package, the regulatory interactions from B where characterized fitting Hill functions to each of the transfer curves, obtaining its characteristic parameters. (D) CcaT+TMA5 inverter dynamics are represented in kymographs. Using the web interface we plotted the growth rate and expression rates for both TUs in time (y-axis) as a function of IPTG concentration (x-axis). Biomass and CFP show a peak at exponential growth and decrease at high levels of IPTG (high dCas9), while YFP peaks at exponential growth as well but increases its expression at high levels of IPTG.

The output of a logic circuit is typically quantified by the reporter expression level,^26,68^ with some threshold determining the output to be high or low. Considering our circuits as logic gates, we quantified their output by the mean expression level. Using Flapjack we can easily plot the output expression level as a function of input IPTG concentration (figure 7B), and use the mean biomass (OD) as a proxy for growth rate. From these induction curves, we can immediately see that growth was strongly affected by induction of dCas9 expression. Further, we observed that both the reference TU and the target TU were repressed in the presence of dCas9. To quantify these regulatory interactions we used the typical Hill function,

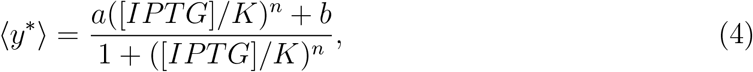

where 〈*y*^∗^〉 is the time-averaged background-corrected fluorescence, *a* is the maximum expression level, *b* is the basal expression level, *K* is the switching concentration of IPTG, and *n* is the Hill coefficicent determining the sharpness of the induction. Using Flapjack this induction curve can be easily computed, plotted, and downloaded (figure 7B) and its corresponding parameter values computed (figure 7C). We compared the Hill function parameters for each fluorescent gene and also computed the Hill function fitted to the mean biomass measurement at each IPTG concentration (figure 7C). This quantity is proportional to the average growth rate over the experiment. The resulting Hill function parameters constitute a datasheet for both on- and off-target regulatory interactions for this set of logical NOT gates or inverters.

Considering these circuits as inverters we must also consider their dynamics. A convenient way to represent time dynamics over a range of inducer concentrations is the kymograph (figure 7D). In these plots each point represents with a color the expression rate at a given time (y-axis) and subject to a given concentration of IPTG (x-axis). To reveal trends the kymographs are normalized to their maximum value. The expression rate of the biomass is the growth rate, and clearly shows a peak at exponential growth and a decrease at high levels of dCas9 (high IPTG). mTurquoise and mVenus expression rates also peak in exponential growth phase consistent with reported growth dependence of gene expression,^69–71^ but decreasing at high dCas9 levels. Surprisingly, the target mVenus gene *increased* its expression rate in stationary phase with high dCas9. This may be due to the dynamics of both dCas9 and sgRNA expression levels (which are not measured). This example illustrates that even circuits encoding simple logic gates can have complex and unpredictable dynamical behaviour that can be characterized using Flapjack.

### Analyzing cell-free TX-TL reactions

Genetic circuits may also be operated *in vitro* using cell-free gene expression, in which transcription and translation reactions are coupled to a cell extract. Gene expression in such systems can be measured in a similar way to *in vivo* assays (figure 8A) using fluorescent reporter genes. Here we use Flapjack to characterize the magnitude and dynamics of four different reporter genes in three different cell extracts.

**Figure 8:**
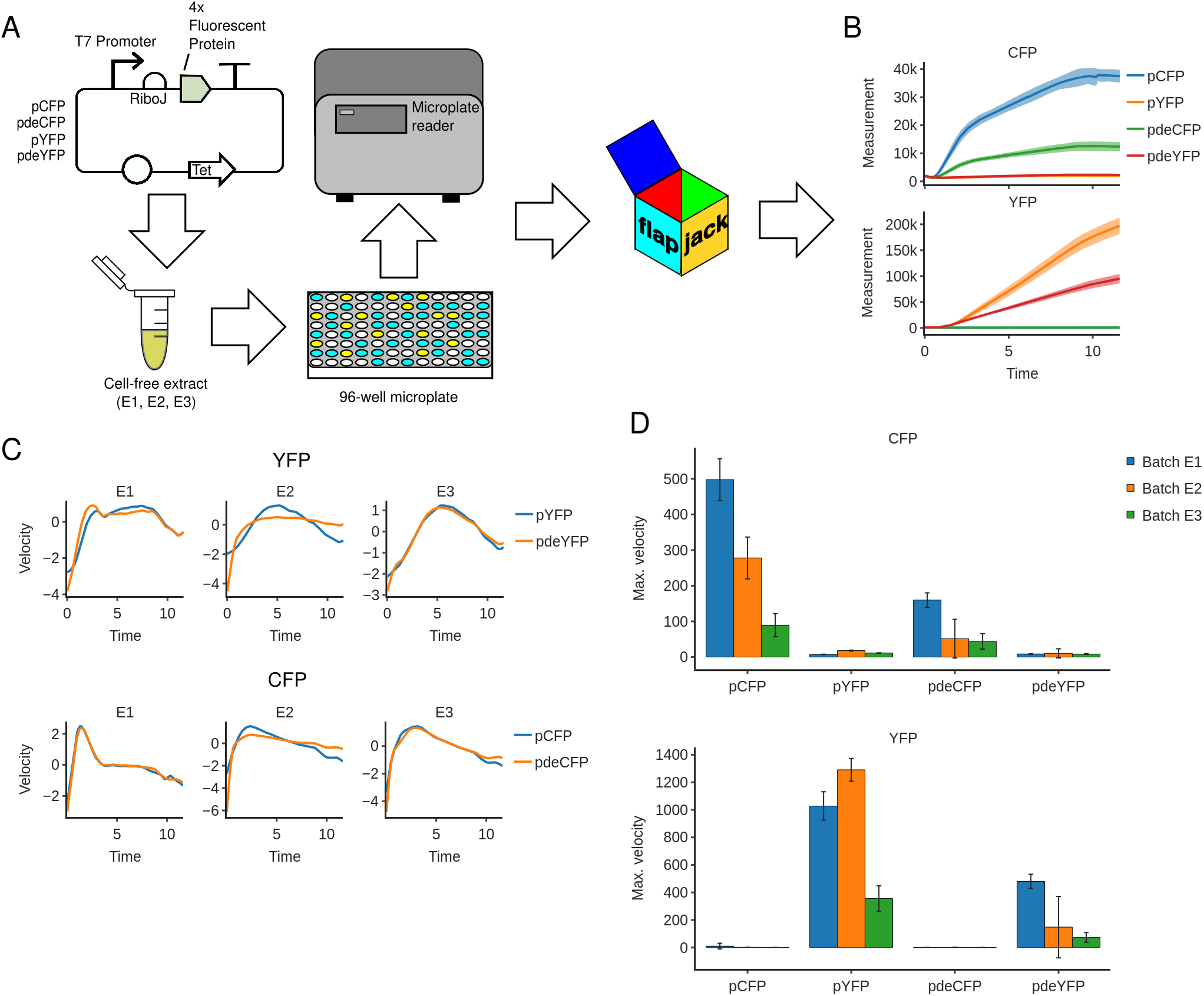
Cell-free fluorescent protein expression. (A) Four different reporter genes (CFP and YFP variants) optimized for transcription and decreased in size, in three different cell extracts. One could expect these variants would less resources and thus produce more fluorescent protein. (B) Counterintuitively, variant reporters did not express more fluorescent protein compared with normal CFP and YFP TUs. (C) Dynamics in cell-free assays cannot be characterized using expression rates, given the lack of a biomass measure. Flapjack describes cell-free dynamics by performing the velocity of protein expression levels. The plasmid coding the fluorescent protein does not affects the shape of the velocity profile, while batches do affect. (D) Finally, and as a complement of the raw expression shown in B, we compute the maximal velocity for each plasmid in all extracts. Again, not optimized TUs show higher velocities than its *de* variants. One can also observe little cross-talk between fluorescent channels, meaning the fluorescence signals were orthogonal.

The reporter proteins were two YFP variants, YFP and deYFP; and two CFP variants, CFP and deCFP.^72–75^ These variants were optimized for transcription and decreased in size such that they have the minimal fluorescence domain. Thus, *de* stands for deletion and enhanced. Given these features, we chose these enhanced and shorter proteins because they may require less amino acids for their synthesis and therefore to reallocate resources of the metabolic load for the synthesis of more protein. Plasmids were constructed so as to constitutively express a single fluorescent protein. In order to test the behaviour of these 4 proteins we prepared 3 cell-free extracts using the protocol described in methods (see Supplementary Information) and incubated with plasmid DNA encoding for a single fluorescent protein (figure 8A). These extracts correspond to different batches of production and allowed us to observe the phenomena of batch-to-batch variability. The genetic circuits were measured in a microplate reader for 12 hours (figure 8A) recording both CFP and YFP fluorescence at regular intervals (figure 8B).

To examine the dynamics of the TX-TL reaction we calculated the reaction velocity over time, defined as *dy/dt* with *y* the fluorescence measurements, for each of the fluorescent proteins in the three extracts. Flapjack calculates the velocity by differentiating the Savitsky-Golay^49^ filtered timeseries. We then normalized each time series by its mean and standard deviation. As shown in figure 8C, the fluorescent protein coded by the plasmid does not affect the shape of the velocity profile. Each batch however shows distinct and consistent dynamics, suggesting that the velocity dynamics are influenced by variations introduced by the cell-free extract preparation protocol. To analyze the intensity of fluorescence expression, we used Flapjack to compute the maximal velocity for each plasmid in all extracts (figure 8D). Firstly we observe that there is little cross-talk between fluorescence channels, meaning the fluorescence signals were orthogonal. Despite the apparently decreased load of the modified *de* fluorescent proteins we note that the original CFP and YFP exhibited higher velocities.

### Image-based measurement and analysis of gene expression

The studies described above were performed using fluorescence microplate readers, however Flapjack and its data model are suitable for any type of kinetic gene expression measurements. Recently low-cost and open source custom-made fluorescence imaging systems have shown that high-quality kinetic gene expression measurements can be obtained^22^ from these alternative sources with potential for high-throughput.^23^ In particular, FluoPi^22^ is a multi-fluorescence imaging station that permits time-lapse experiments at scales ranging from single colonies to entire petri plates. It is low cost and small, enabling high throughput measurement via multiple devices running in parallel, and can be used to perform assays ranging from microbiology to ecology. FluoPi uses low-cost filters that can accomodate excitation and emission wavelengths of long Stokes-shift fluorophores.^22^ In order to use the system it is therefore essential to characterize the behavior of reporter proteins in the device and understand how they relate to image intensities.

These images were analyzed to extract colony size and fluorescence traces that can be analyzed using Flapjack. We used JSON to encode and upload the results of analyses of time-lapse image data from the FluoPi Python package to Flapjack. We analyzed three fluorescent proteins - CyOFP1, mBeRFP and sfGFP - each of which was transformed into E. Coli Top10 strain and grown in a petri dish inside a FluoPi device for approximately 50 hours (figure 9A). We manually selected colonies containing each of these reporters and then used the FluoPi Python package to extract the colony areas and the signal intensities in the red, green and blue camera channels, which were exported to JSON and uploaded to Flapjack for analysis (figure 9B). As seen in figure 9B, colonies have a low level of biomass at the beginning of the experiment, the indirect method for computing the expression rate is not appropriate and thus we use the direct method instead (see above). We computed the mean growth rate using the direct method for each of the three plasmids, pCyOFP, pmBeRFP, and psfGFP, normalized to mean and standard deviation. The plasmids appear to have no effect on growth rate, with colonies on average reaching peak growth at around 24 hours (figure 9C).

**Figure 9:**
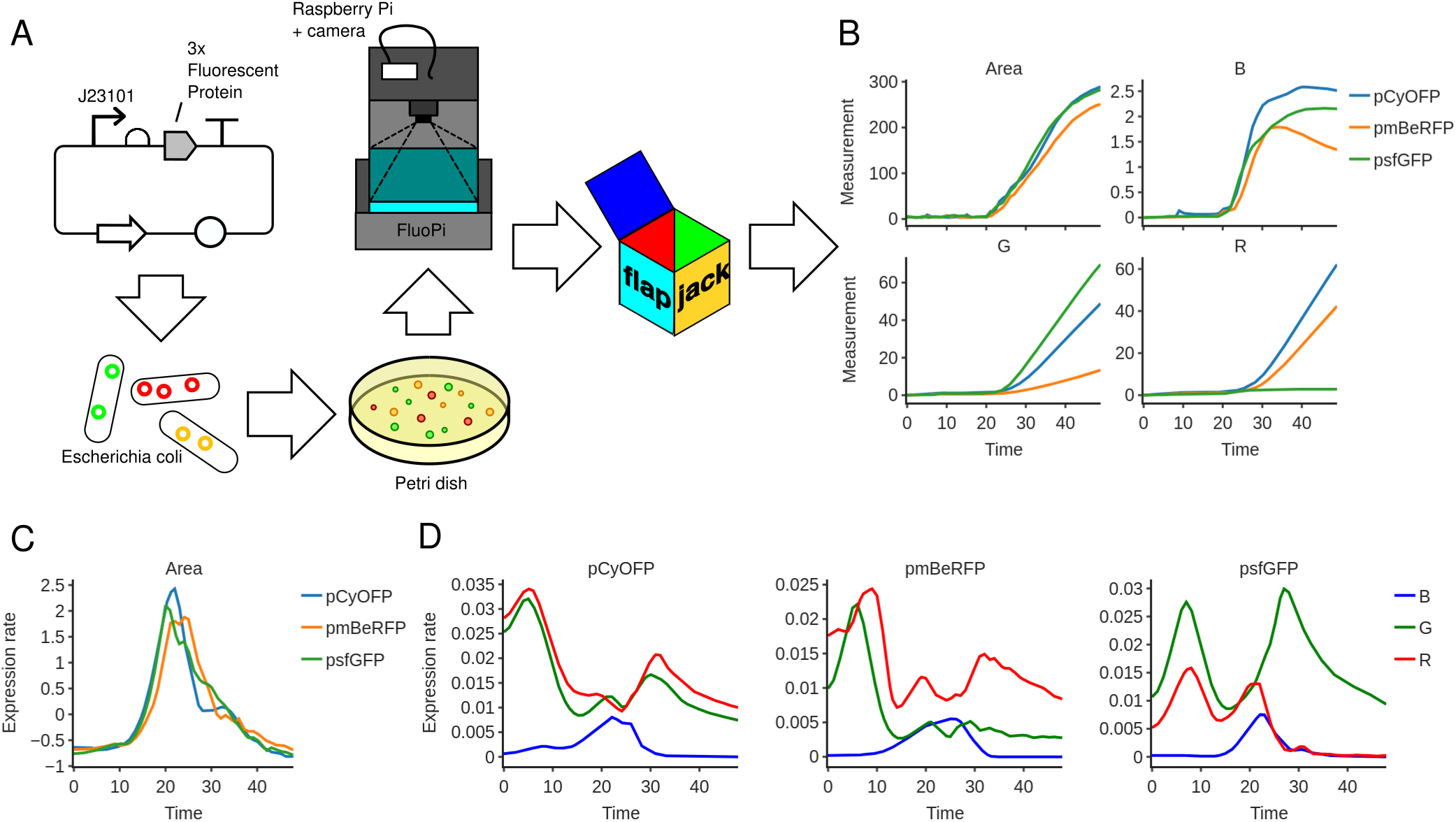
FluoPi imaging station. (A) Plasmids encoding the best 3 fluorescent reporters selected to be used in FluoPi. (B) Using FluoPi Python package, we extracted the colony areas and the signal intensities in R, G and B camera channels from hand-selected colonies. In order to know which expression method to use (direct or indirect) we explore the data, noting that there is low biomass levels. (C) Given these low biomass levels, it is more convenient to use the direct method for computing the growth rate. Plasmids appear to have no effect on growth rate, with colonies reaching peak growth at around 24 hours. (D) Same as for growth rate, expression rates were computed using the direct method. All expression rate profiles exhibit complex double-peaked dynamics for green and red channels, while blue channel show a single peak around 24 hours, suggesting a dependence on growth rate.

We then examined the dynamics of gene expression via the direct linear inversion method applied to each camera color channel (figure 9D). Since the filters of the camera do not correspond directly to the wavelengths of the fluorophores we expect to see bleed-through. Indeed for CyOFP (an orange fluorophore) we see that the fluorescence is recorded equally in the red and green channels. Interestingly the dynamics of each channel are different for mBeRFP and sfGFP, which we expect to be dominated by red and green channels respectively. All expression rate profiles exhibit complex double-peaked dynamics. The blue channel peaks around peak growth suggesting a growth-rate dependence of autofluorescence of cells.^76^ These data show that care must be taken to interpret the results of multi-channel imaging systems, and that more complex analysis may be required to extract the true gene expression profiles.

## Discussion

Synthetic biology aims to apply the engineering principle of design, build, test and learn to biological systems, either to mimic existing functions in nature or to create novel genetic components, networks and pathways. The engineering cycle relies heavily on characterization of genetic parts, devices and systems, creating a need for analysis tools that integrate experimental data with DNA sequences. Recent developments in high-throughput measurements and robotics automation can generate large volumes of data. These data must also be linked to metadata or labels that give context to the measurements, such as growth conditions, host strain, and supplementary inducer chemical concentrations. We developed a system that enables scaleable data storage, analysis, visualization and parameter estimation, enabling collation and sharing of data from diverse sources and different formats. Our datamodel links measurements to metadata describing the conditions under which they were taken, and to the design of the genetic circuit under study through connecting directly to an existing ecosystem of standards and software tools.

We demonstrated that Flapjack closes the loop on the DBTL cycle’s build and learn phases by the use of interconnected tools and modern architecture. The platform makes use of the latest technologies in web development. Its microservices architecture via the use of Docker virtualizations makes installation and deployment easier for users and developers. Flapjack’s backend engine was developed using Django and Django Rest Framework, thus supporting many Python tools especially for data analysis. Despite their usefulness, user experience is key to gain tools usability and increase the community of users. We have made use of frameworks such as React, Redux and D3.js to ensure a fast interface that bridges our powerful storage and analysis engines in an efficient and user-friendly fashion.

Our system is based on an intuitive and accessible web interface, and a powerful API that can be accessed programatically. This could significantly decrease time consuming and error prone data wrangling, allowing characterization of genetic circuits that seamlessly incorporates new data for reliable parameter estimation. It could significantly enhance data sharing, management and analysis for synthetic and systems biology. Given Flapjack’s ability to handle huge amounts of data, it leverages the resources to develop machine learning algorithms and automation protocols.

## Methods

The web application was developed using a microservices architecture with Docker containers virtualization system. For the frontend the React framework was used for the components development, and Redux for state management. For visualizations the library D3.js was used via Plotly. The backend uses Django framework and Django Rest Framework for the API development. WebSockets is used for functionalities that require persistent connection. Persistent data is stored in a PostgreSQL database, while WebSockets data is stored in a Redis database. JSON web tokens are used for authentication. NGINX is used a reverse proxy.

All experiments were performed using standard molecular biology protocols, and along with microplate reader and imaging station assays are described in detail in Supplementary Information.

## Acknowledgement

GYF was supported by Beca Ayudante Doctorando scholarship from the Department of Chemical and Bioprocess Engineering, Pontificia Universidad Católica de Chile. GV was supported by a scholarship from the Institute for Biological and Medical Engineering, Pontificia Universidad Católica de Chile. TJR, GYF, GV, MMS were supported by ANID PIA Anillo ACT192015. FF, AA, IN, TM were funded by ANID - Millennium Science Initiative Program - ICN17 022, Fondo de Desarrollo de Areas Prioritarias - Center for Genome Regulation (ANID/FONDAP/15090007) and CONICYT Fondecyt Iniciacion 11140776. INN was supported by VRI grant from Escuela de Graduados de la Vicerrectoría de Investigación UC. AAM was supported by Beca Doctorado Nacional Conicyt 21140714 - Millennium Science Initiative Program - Millennium Institute for Integrative Biology (iBio). TJR, GYF, GV, MMS were supported by CONICYT Fondecyt Iniciacion 11161046. The authors thank Alvaro Olivera for his support with the iGEM kit.

## Supporting Information Available

Experimental procedures and characterization data for all new compounds can be found in the Supplementary Information document.

## Author Information

### Author Contributions

TJR and GYF designed the study. GYF, TJR, BEG and VCB wrote the software. All authors contributed to writing the manuscript. MMS, INN, TFM, AAM and JD performed the experiments.

### Notes

The authors declare no competing financial interest.

