## Supplementary material for "Flapjack: a data management and analysis tool for genetic circuit characterization"

Table 1

| Symbol | Promoter |
|---|---|
| A | J23101 |
| B | J23106 |
| C | J23107 |
| D | R0011 |
| E | R0040 |
| F | pLas81 |
| G | pLux76 |

Table 2

| Reporter | RBS | CDS | Terminator |
|---|---|---|---|
| RFP | BCD2 | mRFP | ECK0818 |
| YFP | BCD12 | EYFP | ECK9600 |
| CFP | B0034 | ECFP | B0015 |

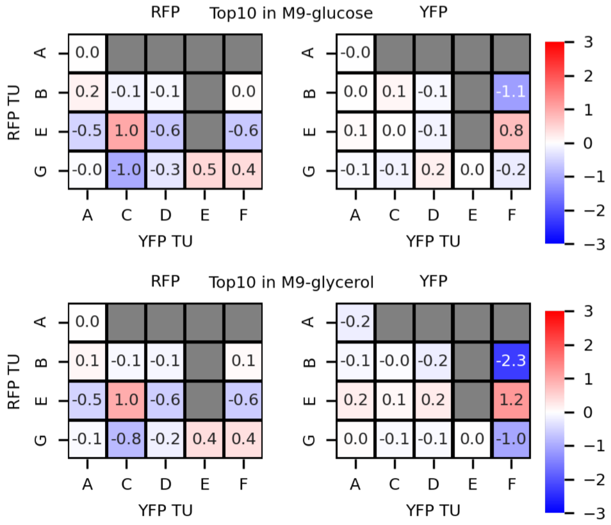


Figure 1: Relative normalized mean expression of TUs in different compositional contexts, for strain TOP10 growing in M9-glucosa and M9-glycerol.

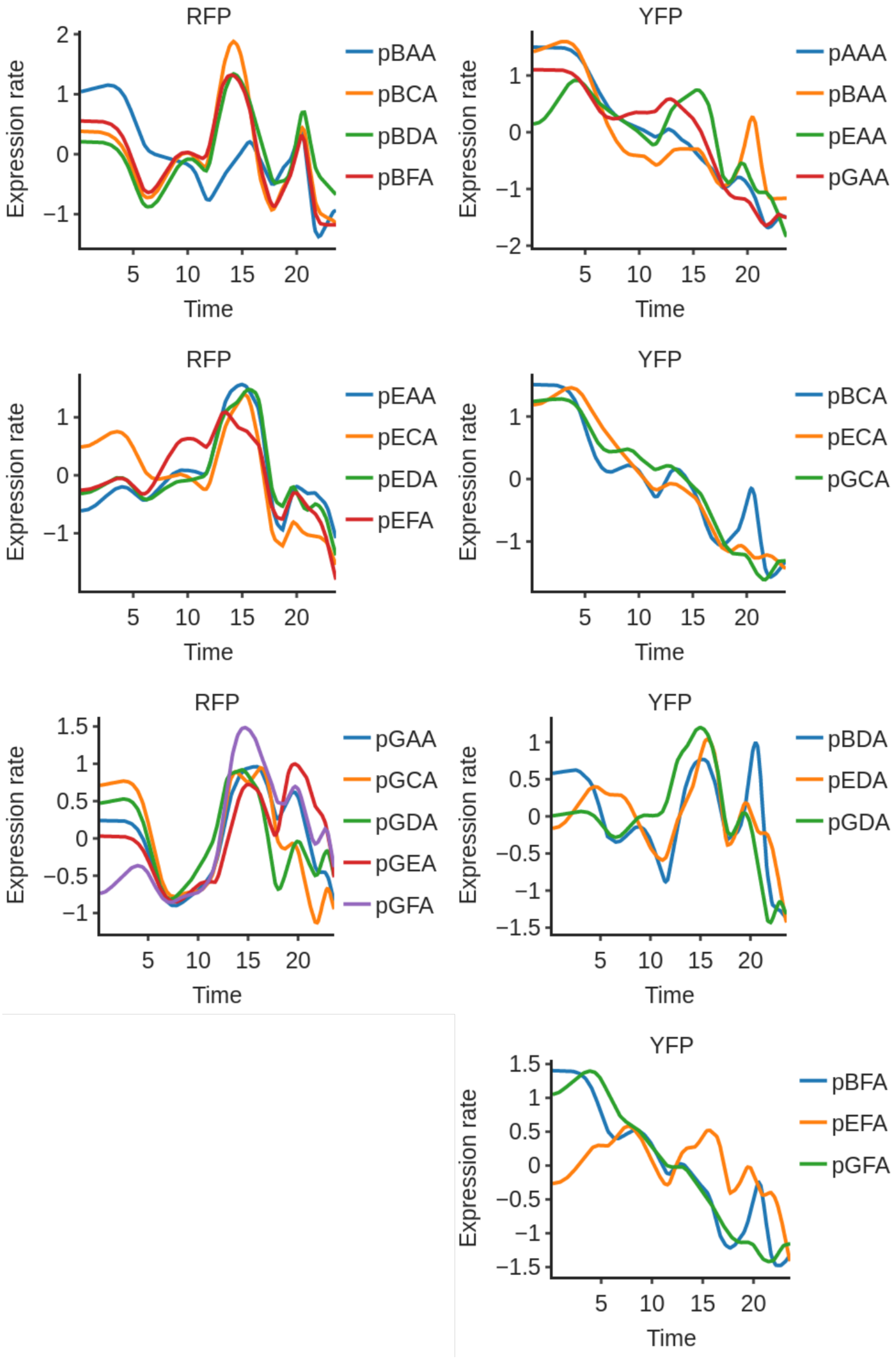


Figure 2: Dynamic gene expression profiles in strain MG1655z1 and media M9-glycerol.

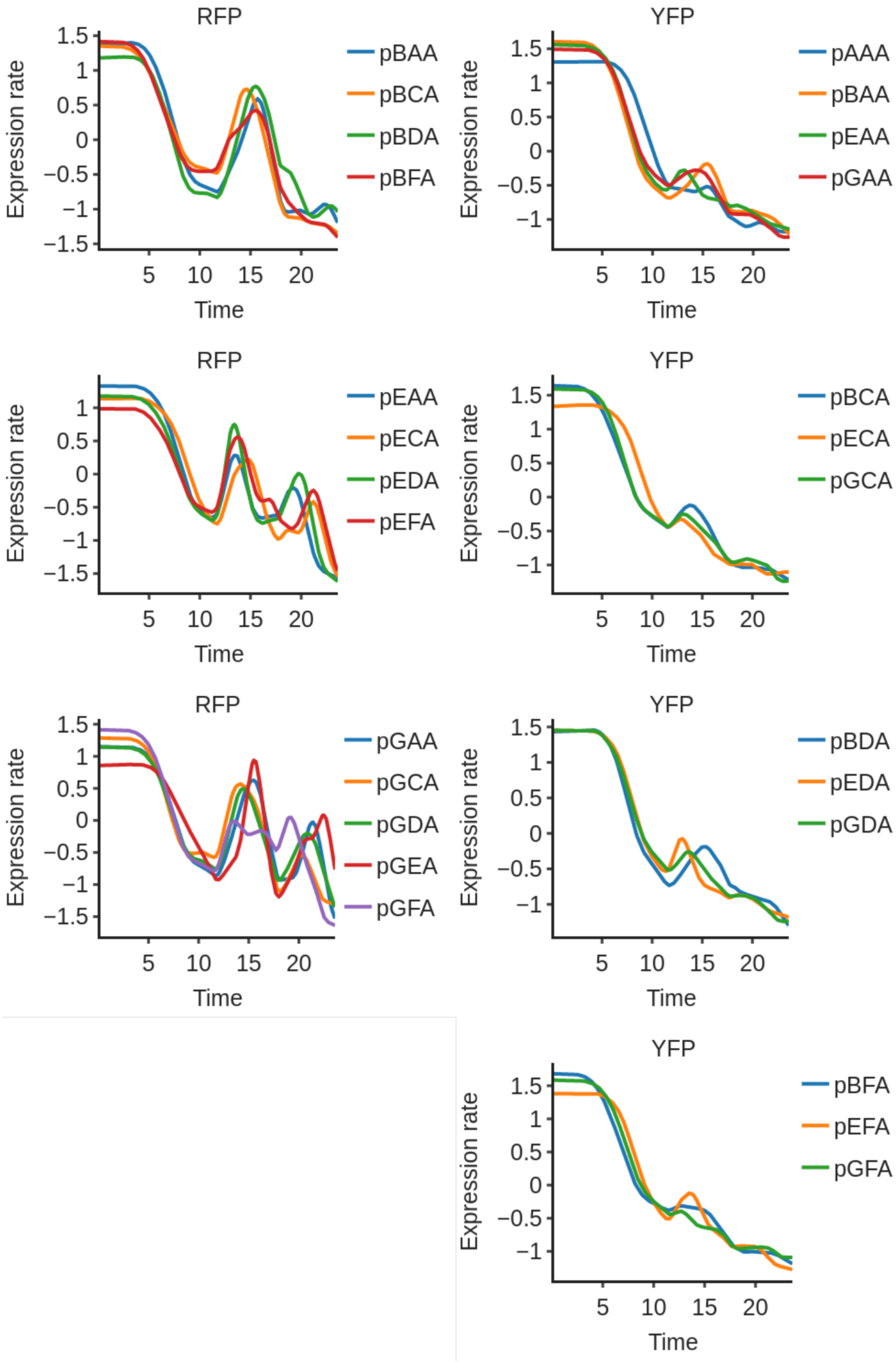


Figure 3: Dynamic gene expression profiles in strain TOP10 and media M9-glycerol.

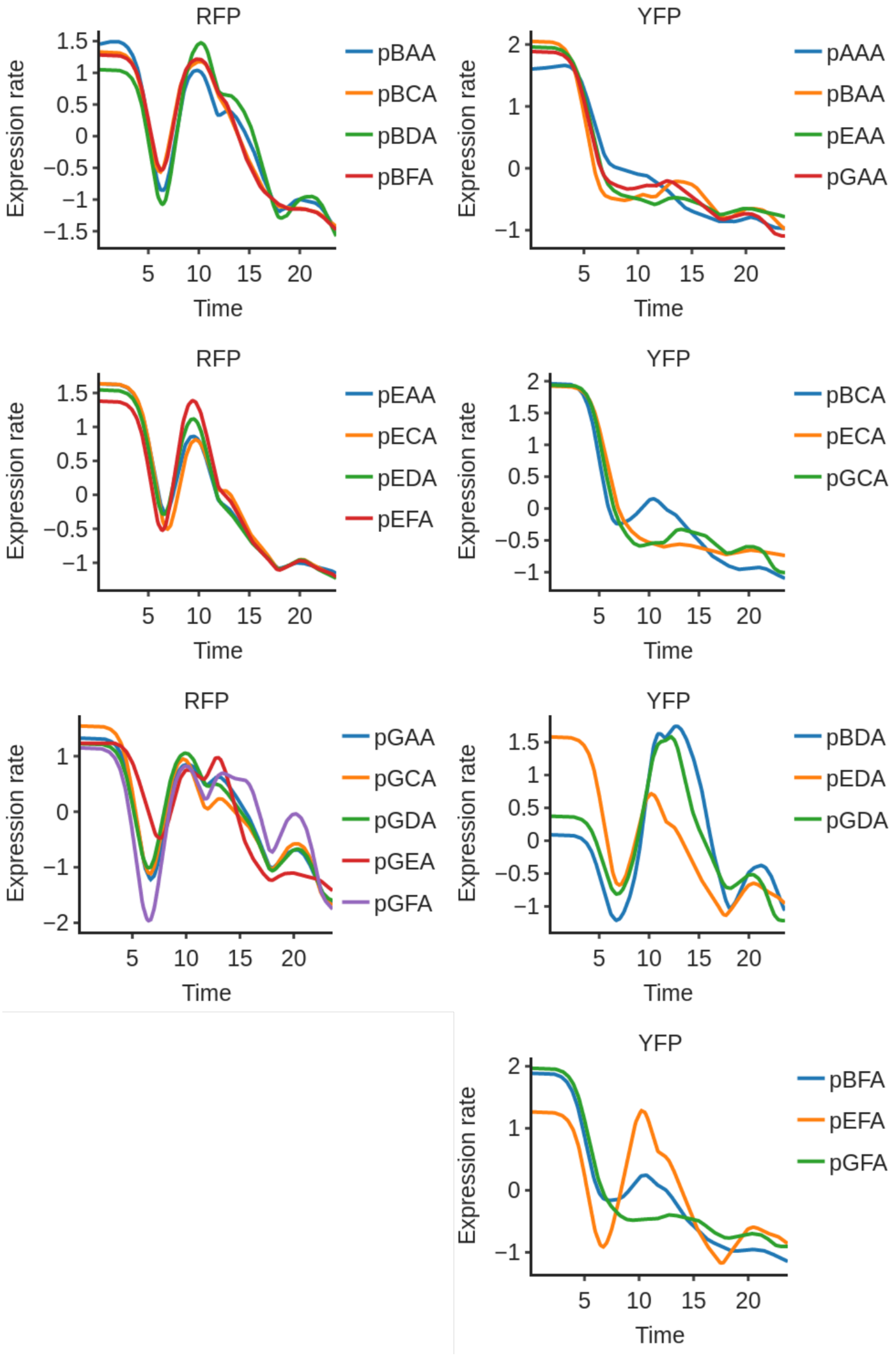


Figure 4: Dynamic gene expression profiles in strain TOP10 and media M9-glucose.

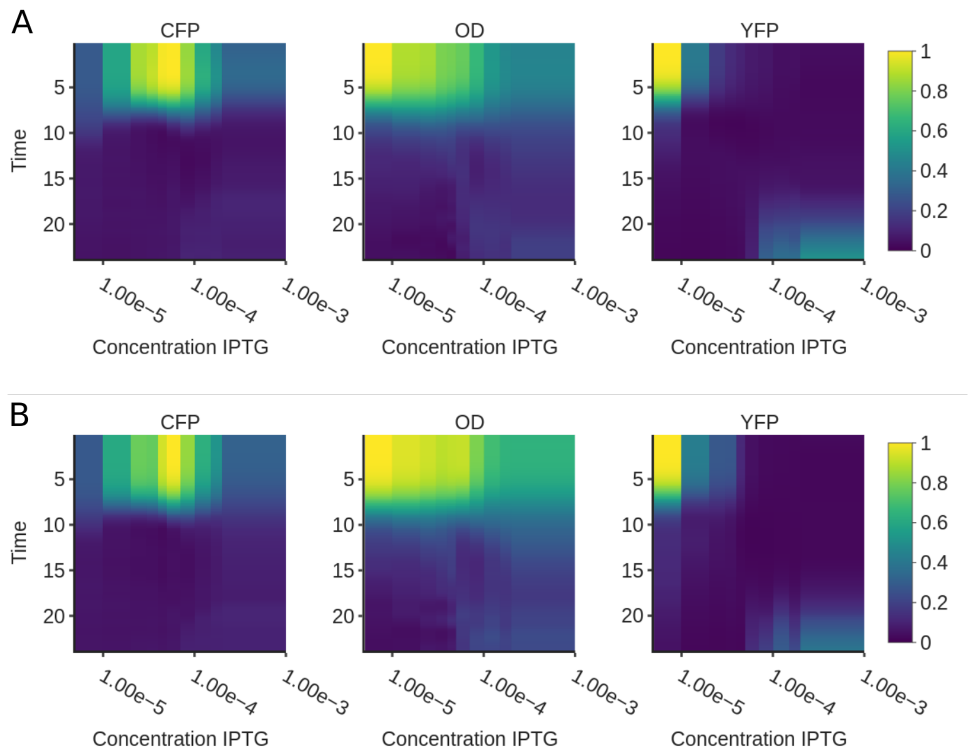


Figure 5: (A) Kymograph for vector CcaT+TMA2.(B) Kymograph for vector CcaT+TMA4.

### Methods

#### Quantifying context effects on genetic circuits

##### Strains

For plasmids selection and stock, the strain used was Escherichia coli TOP10 chemocompetents cells. For making growth assays, the strain used was Escherichia coli MG1655Z1 malE, which has constitutive levels of LacI and TetR repressors.

##### Plasmids assembly

Ten transcriptional Units (TU) were built by Golden Gate Assembly using level 0 DNA parts, consisting of seven Promoters, two Ribosome Binding Sites (RBS), three Coding Sequences (CDS), two terminators and two acceptor plasmids. For each TU, a mix of the level 0 parts was made, calculating equimolar amounts of these, obtaining 20 fmol of each part and 10 fmol of acceptor plasmid in a final volume of 10 $\mu$l. From this mix, 7 $\mu$l were used for the assembly reaction, adding 1$\mu$l of T4 DNA Ligase Buffer 10X (NEB), 1$\mu$l of T4 DNA Ligase 20 U/$\mu$l (NEB) and 1 $\mu$l of BsaI 10 U/$\mu$l (NEB),each reaction had a final volume of 10 $\mu$l.

Fourteen three-reporter plasmids were built by Gibson Assembly. For this, TUs were amplified by PCR to obtain them as linear fragments to be inserted in a destination vector (1X_p15a_Cyan) built previously by the same method. For each plasmid Gibson reaction, a mix of of 4 $\mu$l of the sprecific linear TUs and destination vector fragments was prepared, using 1.5 $\mu$l to mix with 4.5 $\mu$l of Gibson Master Mix, and incubated in a thermocycler at 50 °C for 60 minutes.

**Plate reader growth assays**

M9 media was used with 0.4% w/v Glucose and 0.2% w/v casaminoacids. For each colony containing a the different three-reporter plasmids optical density and fluorescence of RFP, YFP and CFP were measured for 24 hours, every 15 minutes, at 37 °C with constant shaking in 96 well black plates. Each growth assay contained 10 replicates of each plasmid and was repeated on 3 different days, it included four wells of non-transformed bacteria and four wells of the M9 media used without bacteria as controls. The measurements were taken with a Synergy HTX plate reader with Gen5 software. The CFP TU was used as a reference following published work.[1,2] Previously to the assays, liquid cultures of the colonies were grown overnight (or 14-15 hours as a top) in M9 media supplemented with 0.4% w/v Glucose and 0.2% w/v casamino acids. The next day 25 ml of fresh M9 media was prepared and distributed in 2 ml tubes, adding 1996 $\mu$l of media, 2 $\mu$l of the corresponding antibiotic and 2 $\mu$l of the bacteria liquid culture, obtaining a final volume of 2 ml. Each colony was prepared in a separated tube. Then 200 $\mu$l of each diluted colony was added to 10 wells (10 replicates), and four wells for controls. Then the plate was closed, sealed and carried to the plate reader.

#### Characterizing CRISPRi transcription regulation

##### Plate reader assays

Top 10 E. coli strains co-transformed with the appropriate plasmids were inoculated from cryogenic glycerol stock in 1 ml fresh M9-glycerol media supplemented with Kanamycin (100 $\mu$g/mL), Carbenicillin (100 $\mu$g/mL) or both in bacterial tubes and grown at 37 °C and 250 RPM for 8 hours. Next, 1:1000 dilutions in M9-glycerol media supplemented with the corresponding antibiotics per assay, plus ATc (216 $\mu$M) in the case of TMAs strains, were prepared. 198 $\mu$l from each of these cultures were loaded on a Nunc flat-bottom 96-well black plate. IPTG solutions at different concentrations were made by serial dilutions of 1000X stock and 2 $\mu$l were loaded on each well. Finally, the plate was sealed with parafilm and was incubated at 37 °C in a Synergy HTX Plate Reader with continuous orbital shaking (282 CPM) for 24 h. OD600 and fluorescence measurements (mVenus ex: 500nm, em: 540nm / mTurquoise2 ex: 420nm, em: 485nm) were taken every 10 minutes. On each plate, two replicates were made for each IPTG level and two whole replicates were performed on two different days.

##### Plasmid and strain elaboration

Plasmid construction was carried out through Gibson Assembly method .[3] CcaT was created by adding a TetR cassette to pAct6 plasmid developed by Nuñez et al. (2016) .[4] The dCas9 coding sequence was PCR amplified from pdCas9 (a gift from Luciano Marraffini, Addgene plasmid 46569). Fluorescence reporters mVenus and mTurquoise2 were commercially synthesized from IDT based on plasmids used in Rudge et al.(2013) .[5] Promoter sequences for TMA2NT, TMA3NT, TMA4T and TMA5T were created by Gibson assembly [3] based on promoter sequences at pAN-PA2-RFP, pAN-PA3-RFP, pAN-PA4-RFP and pAN-PA5-RFP vectors from Nielsen et al., (2014) .[6] A riboinsulator RiboJ from Nielsen et al., (2016) [7] was used upstream mVenus and mTurquoise2. TMA plasmid backbones correspond to pAct vector backbones used in Nuñez et al. (2016) ,[4] low copy plasmids (pSC101*, Lou et al., 2012 [8]) containing UNS sites for Gibson combinatorial assembly of transcriptional units.

In each case, 5 $\mu$l of the reaction was transformed into CCMB80 chemo-competent E. coli Top 10 cells and plated on LB media with proper antibiotic (Kanamycin 50 $\mu$g/ml, Carbenicillin 100 $\mu$g/ml). Two colonies were selected and propagated to perform plasmid extraction with Wizard® Plus SV Minipreps DNA Purification System (Promega) and checked by sequencing (Macrogen). Co-transformed E.coli strains were obtained by the same transformation method but adding 100 $\eta$g of each plasmid at once. After recuperation they were plated on LB with both antibiotics (Kanamycin 50 $\mu$g/ml and Carbenicillin 100 $\mu$g/ml)

#### Analyzing cell-free TX-TL reactions

##### Preparation of cell-extracts

Cell-extracts were performed similarly to a previously described protocol ,[9] with the following modifications. The cell-extracts were prepared by cultivating E. coli strain BL21 (DE3) in 2xYTPG media (10 g/L yeast extract, 20 g/L Tryptone, 62 mM Na2HPO4 (pH 7.0), and 100 mM glucose) at 37 °C at 300 RPM. The cultures were inoculated at OD600 0.1, induced with 1 mM ITPG at OD600 0.6, and harvested around OD600  4.5. The cells were washed three times in buffer B (10 mM Tris–acetate buffer (pH 8.2), 14 mM magnesium acetate, 60 mM potassium glutamate, and 1 mM dithiothreitol (DTT)). Pellets were weighed and stored at -80 °C for later use. Cells were thawed on ice and resuspended in 1 ml buffer B per g cell wet weight and lysed by sonication with the cell suspension tube in an ice water bath to prevent excessive heating. The lysed cells were centrifuged at 12000 g for 10 min, the supernatant was split into aliquots and stored at -80 °C for later use. All chemicals were provided by Sigma-Aldrich.

##### Cell-free protein synthesis reactions

The protocol for the cell-free protein synthesis was modified from a previously published method .[10] For 1 ml reaction, the following was mixed; 500 $\mu$l cell extract, 20 µl 50xF-mix (1 M K2HPO4), 250 $\mu$l 4xT-mix (960 mM HEPES pH 8.2, 686 mM potassium glutamate, 8% PEG8000, 19.1 mM magnesium chloride, 240 mM glucose, 256 $\mu$g/ml folinic acid, 4.8 mM ATP, 4.8 mM GTP, 3.4 mM CTP, 3.4 mM UTP, 16 mM Cysteine, 8,2 mM each of the other 19 amino acids), 6.7 $\eta$g/$\mu$l DNA, H2O to 1 ml. Individual reactions of 5 $\mu$L of cell-free containing 6$\eta$M DNA were prepared on ice and placed in individual wells in a 96 well plate with V-bottom and sealed immediately to prevent evaporation. Once the plate was loaded, it was placed at 24 °C in a Synergy HTX plate reader (BioTek) and fluorescence (YFP: 540/27 nm excitation, 540/25 nm emission; CFP: 420/50 nm excitation, 485/20 nm emission) was measured every 2:36 min for 12 hours.

##### Plasmid construction

All plasmids shown in this work correspond to Level 1 (single TU) plasmids that were constructed by Golden Gate assembly from Level 0 parts. The level 0 parts were previously prepared by Gibson assembly .[3] All plasmids were sequenced before use.
